# Laurasian legacies in the Gondwanan tree fern order Cyatheales

**DOI:** 10.1101/2023.02.13.528358

**Authors:** Santiago Ramírez-Barahona

## Abstract

Present-day geographic and phylogentic patterns often reflect the Gondwana–Laurasia separation and subsequent history of continental drift. However, some lineages show non-overlapping fossil distributions relative to extant species and in some cases extant ‘Gondwanan’ lineages have ‘Laurasian’ extinct relatives. Here, I combined distribution data for 101 fossils and 442 extant species of tree ferns (Cyatheales) to reconstruct their biogeographic history over the last 220 million years. The time calibrated tree showed most tree ferns families originating during the Jurassic and the onset of crown group diversification beginning during the Cretaceous; a major shift in diversification rates occurred in the largest tree fern family (Cyatheaceae) that comprises ~90% of extant diversity in the group. Biogeographical reconstructions based on extant distributions alone supported a Gondwanan origin for the group; the most probable ancestral range encompassed Australasia and South America. Alternatively, incorporating fossil distribution data into reconstructions showed a Laurasian origin and a most probable ancestral range in Eurasia. These results evince the Laurasian legacies of the Cyatheales spanning the Triassic–Cretaceous, which otherwise remain hidden from biogeographic inferences. These results show that extant-only biogeographic analyses are limited when fossils distribution are more wide spread than in the present-day, highlighting the need to directly incorporate fossils into biogeographical analyses and improve the reliability of ancestral geographic range estimation.

## Introduction

The present-day patterns of species distributions have been shaped by complex biogeographic processes, such as long-distance dispersal, range fragmentation, and local (regional) extinction^1,2^. These processes are interrelated with the geological and climatic history of the planet and observed distribution patterns of organisms often reflect this history. A prominent example of this earth–biota interrelation involves the breakup of the supercontinent Pangea over the last 250 million years^3^. The present-day geographic and phylogentic patterns of animal and plant diversity^4–6^ evince the initial North-South separation into Gondwana and Laurasia that started in the Late Jurassic (~250–200 million years ago)^3^.

Based on the distribution of extant species and tectonic models, reconstructions of clades’ biogeographic histories have given credit to the Gondwanan or Laurasian legacies in distinct lineages^7–10^. At their core, these biogeographical analyses assume that neontological data alone is sufficient to unravel a lineage’s biogeographic history^1,2,11^. However, extinct members of several lineages often show non-overlapping distribution ranges relative to that of their extant relatives^12–15^. This is particularly evident for old Gondwanan lineages, such as ferns, that inhabited the ancient tropical climates of Antarctica^16–18^ and the Northern hemisphere^19^. Non-overlapping distribution patterns can be found in modern-day families across several fern orders (e.g., Gleicheniales^16,20^, Marattiales^21,22^, Salviniales^23,24^, Schizeales^25^, Polypodiales^26^, and Cyatheales^18,27–31^), whose origin most likely predates the breakup of Pangea^32^, leading to wide spread global distributions through time^33^. Several of these fern lineages likely experienced pervasive and geographically heterogeneous extinction rates and therefore reconstructing their biogeographic history based on neontological data alone can result in incomplete and biased inferences^2,10,11,13,34^.

Integrating information of past extinction into event-based biogeographical reconstructions is highly relevant^2,35^ and recent strategies and models have been developed for this purpose^10,11,35^. Few studies have directly incorporated distribution data of fossils to address the biogeographic history of lineages^6,10,11,13,34,36^, partly due to the challenges imposed by the incomplete and fragmentary nature of the fossil record. As with any ancestral state reconstruction, biogeographic inferences are bound to the observed state-space (in this case the present-day geographic distribution of species)^37^; in the presence of extinct unsampled lineages^35^, these limitations can compromise ancestral range estimation. In such cases, biogeographic reconstructions can lead to paradoxical situations where the geographic distribution of the oldest known fossil of a clade (often used to calibrate the age of a lineage) is inconsistent with inferred ancestral ranges^36,38^. These discrepancies highlight the need for an integration of paleontological and neontological data into models of range evolution, particularly in old and wide spread lineages.

The tree fern order Cyatheales is the second largest fern order (~750 species), having a global distribution and an estimated age of origin of 210–230 million years^32,39^. Currently, the eight families in the order are mostly distributed in tropical and subtropical regions^38,40,41^ and less than 1% of species extend into temperate northern regions; however, the group has a rich fossil history showing high species diversity and a wider distribution during most of the Jurassic and Cretaceous. Biogeographic reconstructions for the two largest families (Cyatheaceae and Dicksoniaceae) have suggested Gondwanan origins and a range evolution reflecting the history of continental drift^7,31,38^. However, the oldest fossils indisputably recognized for these families have been found in North America and Siberia^18,27,29,38,42^, well outside their extant (or inferred ancestral) range. Non-overlapping distribution patterns can also be observed for other tree fern families, whose extinct members inhabited regions presently devoid of their extant relatives (e.g., Culcitaceae^30^, Loxomataceae^43^, Thyrsopteridaceae^44^). The presence of Laurasian Cyatheales is not necessarily incompatible with a Gondwanan origin, yet their existence suggests a more complex biogeographic history likely involving pervasive regional extinctions.

To properly assess the biogeographic history of this fern lineage, here I employed models of range evolution including information on the distribution of extant and extinct members. More specifically, I assess the Laurasian legacies of the Cyatheales by estimating divergence times and reconstructing their biogeographic history over the last 230 million years using distribution data on 101 fossils and 442 extant species; I explore heterogeneity in diversification rates across the order within the context of their inferred biogeographic history. These reconstructions show how inferences on dispersal and extinction events, and ancestral ranges can be reshaped when confronted with information from the fossil record.

## Materials and Methods

### Fossils and (paleo)distribution data

I compiled a set of 101 tree fern fossil specimens, including fossilized stems, fertile fronds, and spores (**Supplementary data 1**). Briefly, I selected the most reliable fossil specimens from a list of 1,432 entries compiled from the Paleobiological Database, which I revised and complemented with fossil reviews^18,45,46^, new fossil discoveries ^30,44,45,47–54^, and commonly used fossils for dating^7,38,55,56^; I excluded fossil specimens for which I could not access the original description. The phylogenetic placement of several fossils, mostly stems, have been previously inferred through phylogenetic analyses^46^, but other fossil assignments rely on the presence of clear apomorphies (for instance, spore morphology) or intuitive criteria. Historically, the Cyatheales have been subjected to changing classification schemes and clade circumscriptions, which translates into uncertainties in fossil affinities; for instance, the name Cyatheaceae has often been used to refer to a clade including most, if not all, of the currently recognized families in the order. Therefore, I revised the original fossil descriptions and defined their affinities based on current taxonomic knowledge^41^.

I gathered distribution data for Cyatheales from the Global Biodiversity Information Facility (GBIF.org, 7 February 2020, GBIF Occurrence Download https://doi.org/10.15468/dl.xxjpoz), which were cleaned following the scripts available at https://github.com/spiritu-santi/Shortfalls. For fossil species, I gather the geographic occurrence data directly from the fossils’ locality information provided in the original descriptions.

### Molecular data, phylogenetic reconstruction, and dating

I assembled a molecular dataset with representatives of the eight families and the thirteen genera recognized for Cyatheales. For this I employed PyPHLAWD^57^, an automatic pipeline that retrieves molecular data from NCBI’s GenBank database, generates ‘clusters’ of likely ortholog sequences using BLAST queries, and aligns each cluster using MAFFT^58^. I used GenBank’s plant database (last updated June 28, 2020) by querying for ‘Cyatheales’ sequences longer than 400 bp, with a minimum sequence identity of 0.2, and a minimum coverage of 0.65. I complemented the resulting alignments with sequence data retrieved from NCBI’s PopSet database (PopSet numbers: 1722035357, 1722034779, 1722034120) and from previously published data^55^ ; I incorporated sequences for additional taxa using Clustal2.0^59^ and AliView^60^ and identified and eliminated species synonyms. The final molecular dataset consisted of DNA sequences for 442 species of Cyatheales (60% of the total diversity) with a total length of 15,249 bp for ten chloroplast regions (protein coding regions: rbcL, atpB, atpA, SQD1, and matK; intergenic spacers: atpB-rbcL, rbcL-accD, trnL-trnF, trnG-trnR, and rps3-rpl16) (**Supplementary data 2**); the per-site missing information was 74% and 30.3% of sites were potentially informative.

I estimated a maximum likelihood phylogeny with RAxML v.8^61^, using the GTRCAT model, automatic bootstrap replicates (autoMRE), and substitution parameters estimated for each marker independently. Sequences for *Blechnum* (Blechnaceae), *Odontosoria* (Lindsaeaceae), and *Hypolepis* (Dennstaedtiaceae) were used as outgroups. I used the resulting ML tree to check for ‘rogue taxa’ and evaluate the overall accuracy of the estimated tree topology; five ‘rogue’ taxa were conspicuously out of place and were removed from the alignment. After removing these taxa I estimated the final ML phylogeny (**Supplementary data 3**).

I reconstructed and dated the phylogeny under the Fossilized Birth-Death process^62^ (FBD) using the 442 extant species and the 101 fossils. Compared to node dating, the FBD process incorporates fossil data directly into the tree-generating model, effectively treating each fossil as an independent leaf in the tree^62^; this eliminates fossil redundancy and allows for the inclusion of more than just the oldest fossil for any given clade. Morphological data can be used within the FBD framework to infer the placement of fossil species in the tree (but see Luo et al.^63^). Due to data deficiency I implemented an FBD without morphological data but with topological constraints imposed on fossil taxa^62,64^. Importantly, I use a single representative for each fossil species in the analyses, with the single exception of *Cyatheacidites annulatus* for which I incorporated two specimens due to its wide spread distribution across space and time.

I implemented the FBD dating in RevBayes^65^ using the following settings (code available at https://github.com/spiritu-santi/biogeograhy): GTR+G nucleotide substitution model with four gamma categories; an uncorrelated exponential clock model; uniform prior in the occurrence time of fossils; and the process conditioned on survival. I used a uniform sampling fraction of 0.60 and set the origin time of the process to 215–230 Mya (estimated divergence times between Cyatheales and Polypodiales^32,39^). To decrease computational burden, I defined several monophyletic constraints on the tree based on well supported phylogenetic relationships^38,41,55,66^. Initially I ran the analyses for 200,000 iterations, but there was poor mixing and several model parameters showed effective sample sizes (ESS) < 100. Thus, I used the final model parameters (that is, branch rates, dated tree, origin time) obtained in this ‘long’ run to set the starting values for two final ‘short’ runs; these consisted of MCMC chains ran for 50,000 iterations each, sampling every 10 iterations, and defining a period of 2,000 iterations to tune the efficiency of the MCMC proposals. To summarize the results, I defined a burn-in fraction of 10% for each run after inspecting the likelihood traces using the ‘coda’ package^67^ in R^68^; most parameters had an ESS > 200 after burn-in. I set up the analyses to produce posterior samples of trees including all fossil as tips and the corresponding trees without fossils. I constructed maximum clade credibility (MCC) topologies with node ages defined from the median node heights as implemented in RevBayes^65^ (**Supplementary data 4–5**).

### Rates of species diversification

I assessed diversification rate variation across the Cyatheales phylogeny by using Bayesian Analysis of Macroevolutionary Mixtures (BAMM) v.2.5^69^ (**Supplementary data 6**). I used BAMM to: (1) test whether a single or multiple macroevolutionary regimes can explain the diversification history of tree ferns; (2) and infer the approximate locations in the phylogeny of shifts in macroevolutionary regimes that are maximally supported by the data. I performed three separate analyses using the MCC topology excluding fossils to assess the influence of per-lineage sampling fractions on the inferred number (and location) of shifts (**Supplementary data 7–8**): (i) a global sampling fraction of 0.6; (ii) a per-genus sampling fraction; and (iii) a per-lineage sampling fraction to account for uneven sampling within genera (for instance, 98% of Old World *Cyathea* vs. 67% of New World *Cyathea*). Priors on rate parameters were scaled to the dated tree using the setBAMMpriors function in the BAMMtools^70^ package in R^68^.

I ran each analyses for 25 million generations, sampling every 1,000 generations, and specifying a burn-in period of 50%. I assessed convergence and effective sample sizes (ESS) using the coda^67^ package in R^68^; in all analyses the likelihood and the number of shifts had ESS > 500. I used the BAMMtools^70^ package in R^68^ to estimate Bayes factors and compare (when possible) the evidence for multiple-shift models against evidence for a model with no rate-shifts^71^. I then visualized the approximate location of shifts by estimating the 95% credible set of shift-configurations using a marginal posterior-to-prior odds ratio of 50 and the maximum shift credibility configuration as implemented in the BAMMtools^70^ package in R^68^.

### Biogeographic reconstructions

I performed ancestral geographic range reconstruction for Cyatheales using the Dispersal-Extirpation-Cladogenesis (DEC) model^72^ with time-dependent effects on dispersal rates as implemented in RevBayes^65^. I defined four connectivity matrices using eight broad areas: (1) North America; (2) South America; (3) Antarctica; (4) Eurasia; (5) Africa; (6) Southeastern Asia; (7) Australasia; and (8) India. The connectivity matrices correspond to four time periods: Triassic–Jurassic (250–145 Mya); Cretaceous (145–66 Mya); Paleogene (66–23 Mya); and Neogene–Quaternary (23–0 Mya). I scored the geographic distribution for extant and fossil species across the eight regions using the geographic occurrence database (code available at https://github.com/spiritu-santi/biogeograhy); in this coding scheme North America includes the Nearctic region and the Northern Neotropics, and Antarctica includes the Southern tip of South America and isolated islands in the Pacific (e.g., Juan Fernandez Island).

To assess the influence of fossil data on the estimation of range evolution, I performed two separate analyses: 1) *extant-only* reconstruction including only living species; and 2) *full* reconstruction including living and fossil species. For each analysis, I ran 5,000–10,000 iterations with sampling of the MCMC chains every 10 iterations to obtain a posterior sample of at least 500; a burn-in period of 200 iterations was established to tune the efficiency of the MCMC proposals. I specified a varying burn-in fraction for each run after inspecting the likelihood traces using the ‘coda’ package^67^ in R^68^. To incorporate uncertainty in tree topology and divergence times, I defined a ‘tree’ parameter in the model by using an empirical distribution drawn from the posterior sample of trees obtained in the FBD dating analyses; this parameter was sampled every iteration based on the regular Metropolis-Hastings acceptance ratio. B doing this, the likelihood estimation of the complete model incorporates, to some extent, the reciprocal effects between the phylogeny and range evolution. Due to computational and data limitations, I did not perform a joint estimation of the dated phylogeny and range evolution^73^. Prior to running the analyses, I filtered the posterior samples of trees to obtain a set of trees with the same family-level backbone topology as that of the ML tree and resolved all sampled ancestors as branches with near-zero branch lengths (0.0001). I constructed maximum clade credibility (MCC) topologies with node ages defined from the median node heights as implemented in RevBayes^65^.

I sampled ancestral states and stochastic mappings for range reconstructions, which were summarized and visualized using MCC topologies, state-frequency-through-time (SFTT) plots, and transition graphs. SFTT plots summarize the proportion of lineages found in different states (areas) at a given time while averaging over the posterior sample of trees and stochastic mappings^73^; I constructed the SFTT plots using 2-Myr bins. Transition graphs summarize the number of times a dispersal event is observed between pairs of areas; in this case, dispersal events take the form Area1 → (Area2 + Area1). Extinction events were summarized by tallying the number of inferred events per area, where an extinction event in Area2 would take the form (Area1 + Area2) → Area1. All plots were constructed using the tidyverse^74^, ggplot2^75^, and igraph^76^ packages in R^68^; plots were further edited using Adobe Illustrator (Adobe, United States).

## Results

### Divergence times and shift in diversification rates

The combined analyses of 101 fossils and 442 extant species resulted in a maximum clade credibility (MCC) tree that agrees with previous phylogenetic hypothesis (**figure 1a**). The dating analyses relied on an estimated stem age for tree ferns of 215–230 Mya, which marks the divergence between Cyatheales and Polypodiales occurring during Late Triassic; accounting for dating uncertainty, the crown node of Cyatheales was dated to 226–188 Mya, placing the age of living lineages in the Late Triassic–Early Jurassic (**figure 1a**). The origins of most tree fern families (stem ages) were estimated mostly in the Jurassic except for the divergence between Plagiogyriaceae and Culcitaceae that occurred during the Late Cretaceous–Paleogene (Supplementary table S1). The age of the onset of crown group diversification of the largest tree fern family, the Cyatheaceae (~90% of extant species in the order), was estimated in the Early Cretaceous (139–105 Mya), whereas the crown group ages of extant genera in the order were estimated to have started in the Late Cretaceous–Paleocene (**figure 1a**).

**Figure 1.**
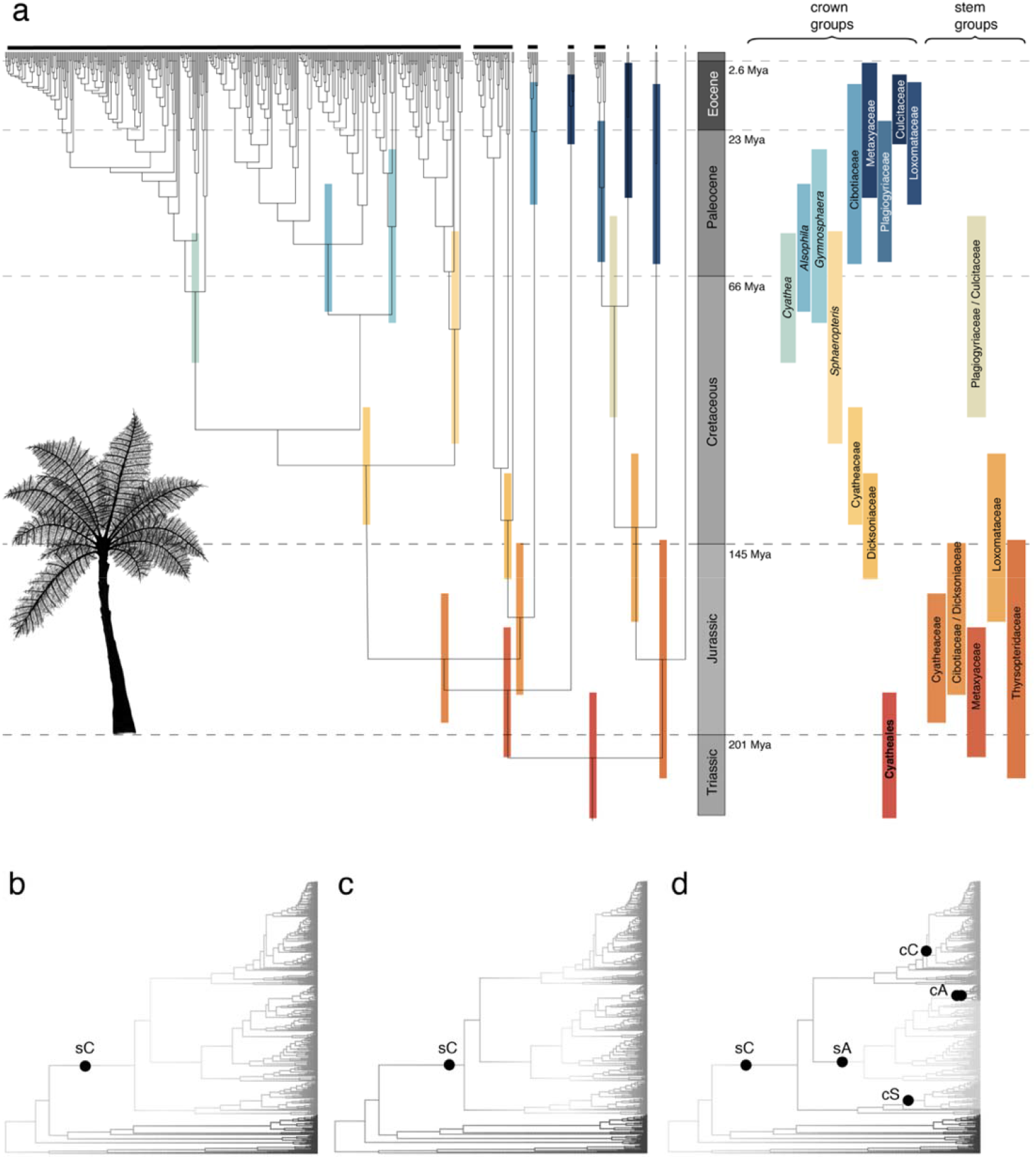
Divergence times and diversification rates shifts in tree ferns (Cyatheales). a, time-calibrated phylogenetic tree of Cyatheales constructed under the Fossilized Birth-Death process showing the 95% HPD for stem and crown nodes of families and genera; the 95% HPD for crown and stem groups are also depicted as horizontal bars. b-d, location of core shifts in diversification rates detected by BAMM across the tree fern phylogeny using (b) global, (c) per genus, and (d) per lineage sampling fractions. Core shifts are named according to their location: sC = stem Cyatheaceae; sA = Stem *Alsophila*; cS = crown *Sphaeropteris*; cC = crown *Cyathea*; cA = crown *Alsophila*.

BAMM detected significant levels of rate heterogeneity across the tree fern phylogeny even under different sampling fractions of extant lineages (**figure 1b–d**). Using a global sampling fraction and a per-genus sampling fraction resulted in best shift configurations with high posterior probabilities (f = 0.62 and 0.77, respectively) supporting scenarios with a single rate shift (**Supplementary figures S1–S2**). Using a global sampling fraction, the 95% credible set included nine shift configurations, out of which only one recovered zero shifts (**Supplementary table S2**).

By implementing a per-genus sampling fraction, the 95% credible set included four shift configurations with one or two shifts (**Supplementary table S2**). Under these analyses, the comparison between the one-shift models against a zero-shift models resulted in Bayes factors > 1000, whereas the comparison between the one-shift models and models with more than one shift resulted in Bayes factors < 1.0 (**Supplementary table S3**). These results gave strong support for a one-shift scenario as the overall best model of rate heterogeneity; the maximum credibility shift configuration showed an acceleration of diversification rates along the stem branch of the scaly tree ferns (Cyatheaceae) (**figure 1b-c**).

The analyses using a per lineage sampling fraction resulted in a 95% credible set that included 655 shift configurations detecting multiple (>3) core shifts in the Cyatheales; the best shift configuration had a low posterior probability (f = 0.08) and recovered four shifts (**Supplementary figure S3**); under this sampling scheme, zero-shifts models were not sampled in the MCMC runs (**Supplementary table S2**). The maximum credibility shift configuration retrieved five shifts in diversification rates, all within the scaly tree ferns (Cyatheaceae), and the shift previously detected along the stem branch of this family (**figure 1d**); two of these shifts occurred along the same branch within *Alsophila* and are therefore depicted as a singly shift (cA) in **figure 1**.

### Biogeographical reconstructions

The biogeographical reconstruction based on the distribution of extant species (extant-only reconstruction, **figure 2**) supported a scenario in which the most recent common ancestor of extant Cyatheales had Gondwanan affinities; the two most probable reconstructed ancestral ranges were Australasia and [Australasia + South America] (**figure 2a-b**). These Gondwanan affinities were shared by the scaly tree ferns (Cyatheaceae), whose reconstructed ancestral range most likely included Australasia (**figure 2b**); other tree ferns families also showed Gondwanan affinities except for the Culcitaceae and the Plagiogyriaceae whose crown groups showed Laurasian ranges (**figure 2b**). The SFTT plot showed a wide spread distribution across Gondwana throughout the last 200 My, initially occupying mostly Australasia during the Early Jurassic and then expanding into other landmasses towards the Late Jurassic–Early Cretaceous (**figure 2a**).

**Figure 2.**
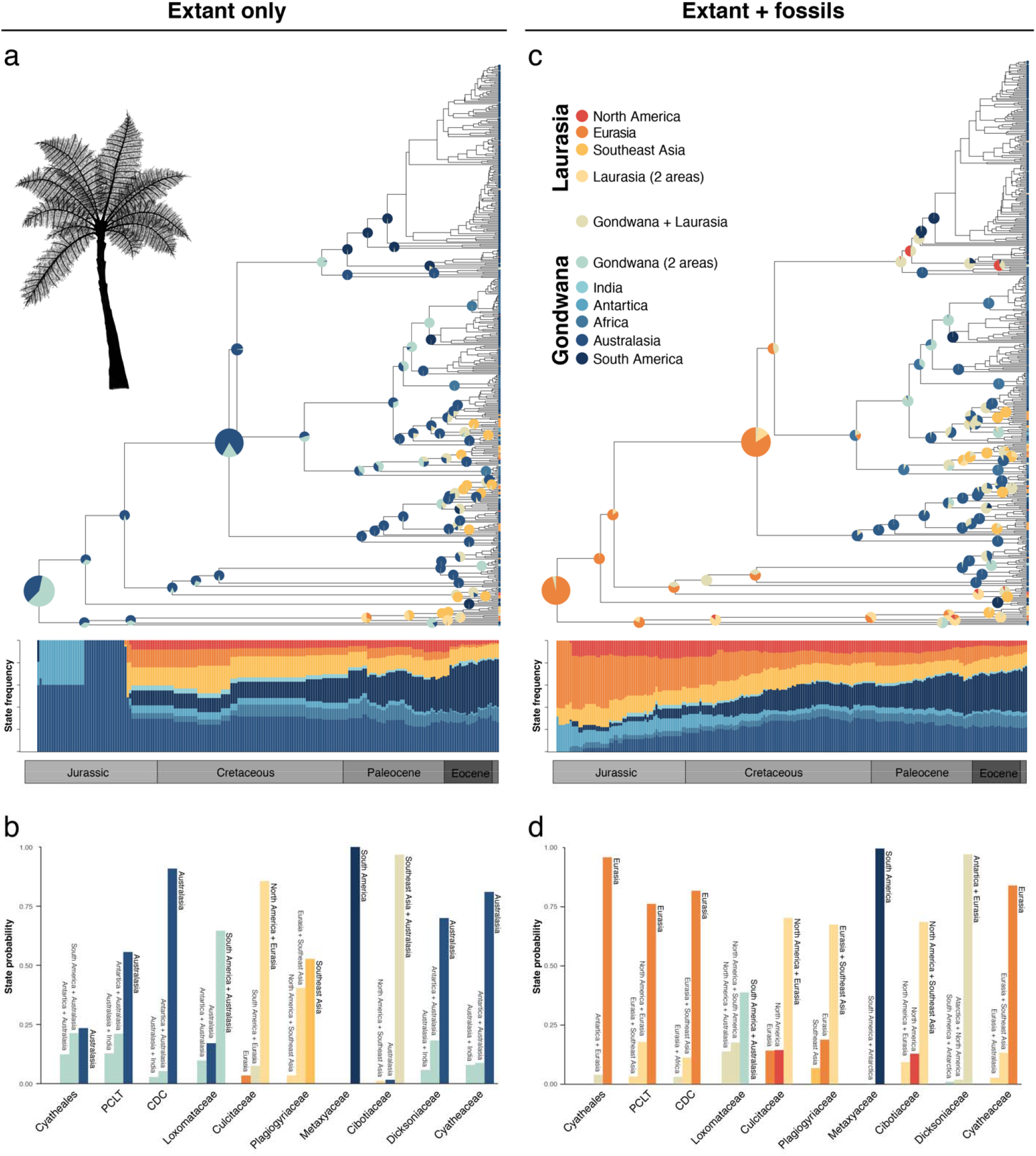
Biogeographic reconstructions for tree ferns (Cyatheales) based on distribution data of extant and fossil species. a, c: phylogenetic trees of Cyatheales showing the most probable inferred ancestral ranges for nodes across the phylogeny based on reconstructions excluding (a, extant only) and including fossils (c, extant + fossils); for simplicity information for several nodes was omitted. State frequency through time (SFTT) plot showing the temporal distribution of probable ancestral geographic states. b, d: posterior probabilities of the three most probable inferred states for each node under reconstructions excluding (b, extant only) and including fossils (d, extant + fossils). CDC represents the node for the most recent common ancestor of the Cyatheaceae, Dicksoniaceae, and Cibotiaceae. PCLT represents the node for the most recent common ancestor of the Plagiogyriaceae, Culcitaceae, Loxomataceae, and Thyrsopteridaceae.

The number of estimated dispersal events showed that range expansions mainly happened out-of-Australasia into the rest of Gondwanan landmasses (**figure 3a**). Dispersal events across Gondwana appeared to be low but constant during the last 200 My, whereas dispersals into Laurasia most probably happened during the Neogene (**figure 3b**). Likewise, South America and Southeast Asia appeared as the source of range expansions of Cyatheales into North America and Eurasia, respectively. There was a higher frequency of regional extinction in Gondwana relative to Laurasia and Australasia had the largest number of regional extinctions (**figure 4a**); extinction events appeared to have been constant in Gondwana during the last 200 Ma but only recently for Laurasia (**figure 4b**).

**Figure 3.**
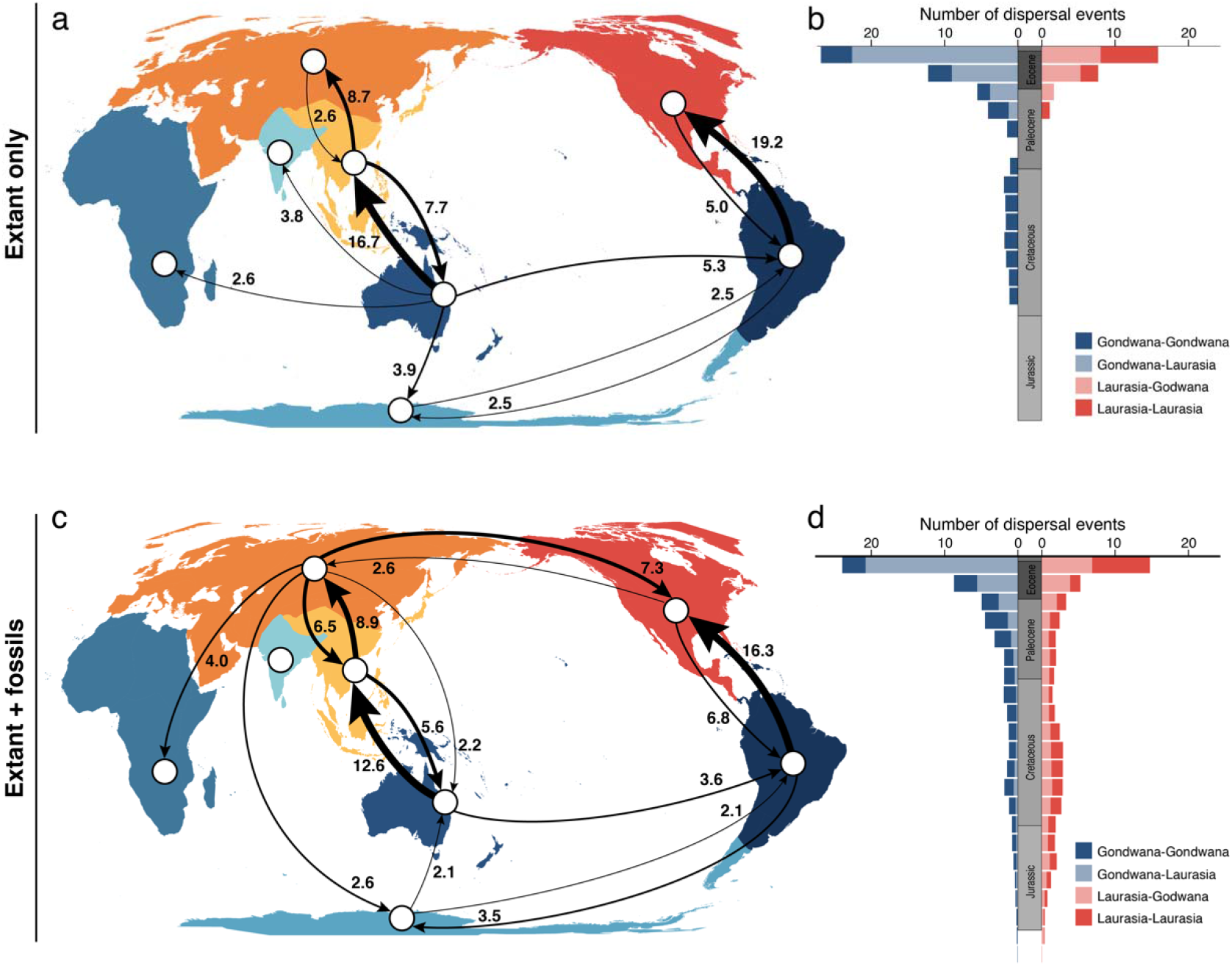
Reconstruction of dispersal events for tree ferns (Cyatheales) across geographic regions based on distribution data of extant and fossil species. a, c: number of inferred dispersal events (posterior mean) among geographic regions based on reconstructions excluding (a, extant only) and including fossils (c, extant + fossils). b, d: number of inferred dispersal events through time based on reconstructions excluding (b, extant only) and including fossils (d, extant + fossils). For simplicity only numbers > 2 are shown in the map. Dispersal event through time were estimated in 10 million year bins and grouped according to the direction of movement among major geographic regions (Gondwana and Laurasia).

**Figure 4.**
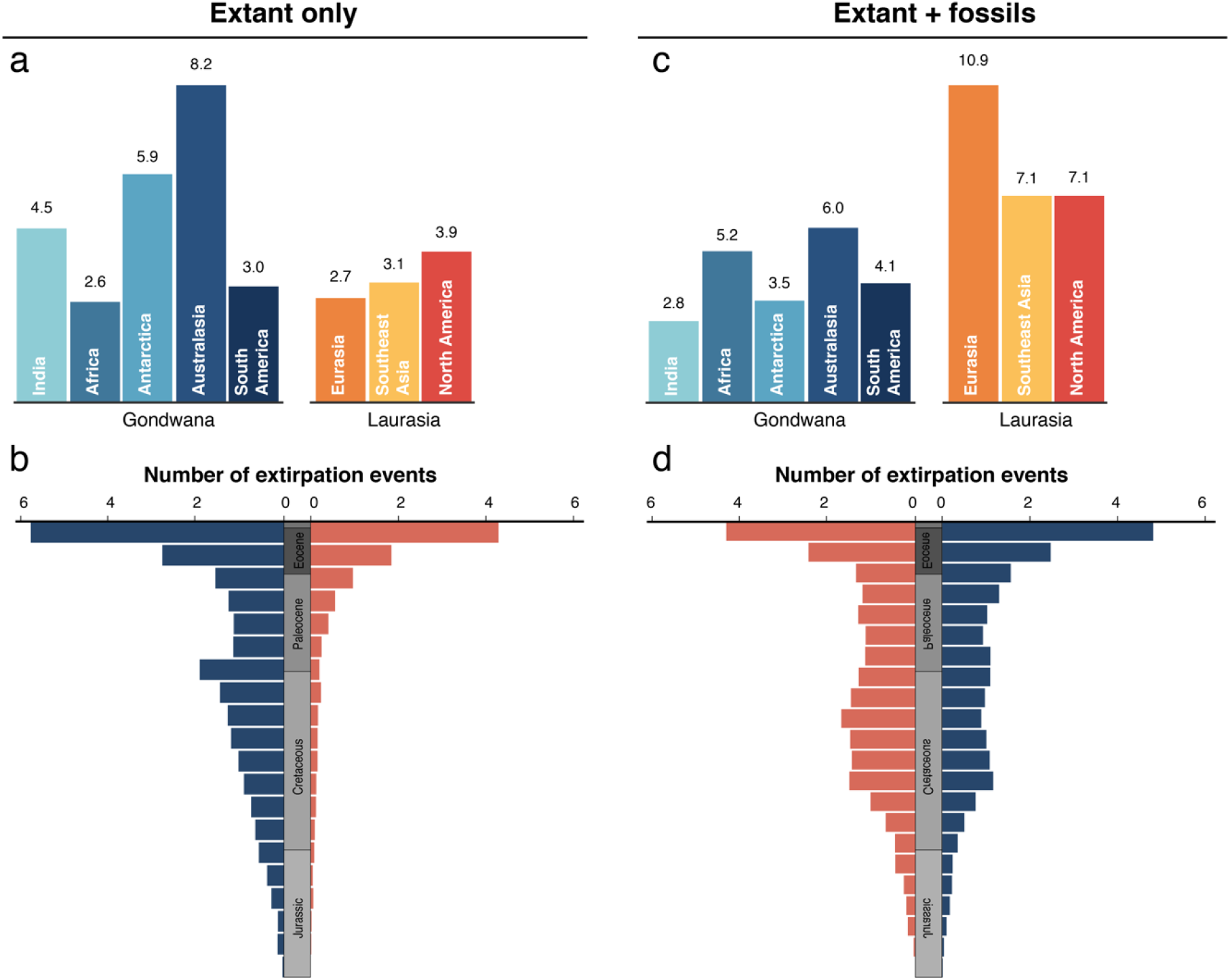
Reconstruction of regional extinction events for tree ferns (Cyatheales) across geographic regions based on distribution data of extant and fossil species. a, c: number of inferred regional extinction events (posterior mean) within geographic regions based on reconstructions excluding (a, extant only) and including fossils (c, extant + fossils). b, d: number of inferred regional extinction through time based on reconstructions excluding (b, extant only) and including fossils (d, extant + fossils). Extinction events through time were estimated in 10 million year bins and grouped according to the major geographic regions (Gondwana and Laurasia).

By incorporating paleodistribution data into the analyses, the analyses provide a radically different alternative of the biogeographic history of the Cyatheales (*full* reconstruction, **figure 2**). This alternative scenario did not support the Gondwanan affinities of crown Cyatheales and instead showed that the most probable reconstructed ancestral range encompassed Eurasia (**figure 2c-d**); this reconstruction retrieved mostly Laurasian affinities of tree fern families expect for the Metaxyaceae and the Loxomataceae (**figure 2d**).

The full reconstruction supported ancient dispersal events into Gondwanan landmasses occurring within crown Cyatheaceae, which most likely occurred during the Cretaceous (**figure 3c-d**). These dispersal events include the range expansion into Gondwana of the four genera of the Cyatheaceae; *Cyathea* retained a reconstructed ancestral range that included Laurasian landmasses until the Paleocene and only latter achieved its current distribution predominantly in South America and a few species in Australasia. Likewise, the most probable ancestral range for stem Dicksoniaceae was Eurasia (**figure 2d**) and dispersal events into Gondwanan landmasses also occurred during the Cretaceous. The SFTT plot showed that Cyatheales had a wide spread distribution across Pangea during the early stages of their evolutionary history and the occupancy of Laurasian landmasses gradually subsided towards the present-day (**figure 2c**). The estimated number of dispersal events showed a high frequency of range expansions from Australasia into Southeastern Asia (**figure 3c**), but a more complex dispersal network was retrieved with a higher frequency of dispersal events involving both Laurasian-Laurasian and Laurasian-Gondwanan range expansions (**figure 3d**). Under the full reconstruction Australasia had the highest frequency of extinction events among Gondwanan landmasses, but slightly lower as those estimated for Laurasian landmasses (**figure 4c**). Extinction events appeared to have been constant during the last 200 My for both Gondwana and Laurasia (**figure 4d**); a slight increase in regional extinctions was observed during the Middle to Late Cretaceous (**figure 4b**).

## Discussion

### Diversification rate heterogeneity in the Cyatheales

The dated phylogeny for the Cyatheales here presented agrees with previously published phylogenies^7,38,55,66^ and provides the most complete account of the timing of evolution across this ancient group of ferns. Stem and crown ages for all families and genera provide evidence of a delayed onset of diversification of extant lineages occurring until the Cretaceous. Previous studies for ferns have provided (stem) age estimates for Cyatheales, but with poor sampling within the order^39,66^. The most recent version of the Fern Tree of Life^32^, dated with a Penalized Likelihood approach, provides similar ages for crown Cyatheales and for the two most speciose families within the order. Studies focused on the Cyatheales have inferred species-level divergence times for the two most speciose tree ferns families^7,38,55,56,77^ that, despite some differences, are broadly consistent with the divergence times here presented.

As observed for other groups of ferns^32^, the divergence times estimated across the Cyatheales suggest that the origin of major living lineages pre-dates the Angiosperm terrestrial revolution occurring towards the Late Cretaceous^78^. However, the inferred divergence times fit the narrative of extant ferns diversifying in the shadow of angiosperms^32,78,79^, suggesting that crown group diversification of Cyathealean genera occurred in synchrony with the rise to ecological dominance of flowering plants during Late Cretaceous–Paleogene. Accordingly, the diversification analyses strongly support a shift in diversification rates along the stem branch of scaly tree fern family (Cyatheaceae) during the Cretaceous and provide evidence of multiple rate shifts likely occurring within this family during the Cenozoic. I argue these shifts are possibly linked to the emergence of modern-day tropical ecosystems during the Paleocene^80,81^; it is within these ecosystems in which most modern-day scaly tree fern diversity is concentrated.

Previous studies have come to contrasting conclusions regarding heterogeneity in diversification rates in the Cyatheaceae^56,82^. Loiseau et al.^56^ suported a history for Cyatheaceae with no significant rate shifts based on BAMM. However, other approaches to estimate rate heterogeneity (BayesRate) revealed significantly higher diversification rates in the most species-rich genera in the family^56,82^, partially agreeing with the present results. Even if rates of diversification have remained constant within the Cyatheaceae^56^, as suggested by two of the three BAMM analyses (**figure 1b-d**), the observed rate shift along the stem branch of the family (during the Jurassic-Cretaceous) likely lead to significantly higher rates of diversification and explain the overwhelmingly higher species diversity of this lineage relative to the rest of the order.

### The biogeography of Cyatheales

The biogeographic reconstruction based on extant species (extant-only) is consistent with the presumed Gondwanan origins of the Cyatheales and agrees with previous studies on the Cyatheaceae^38^ and the Dicksoniaceae^7^. The SFTT plots show that Cyatheales were initially wide spread across Southern Pangea (later Gondwana) during most of the Jurassic and subsequently occupied Laurasia via secondary dispersal. Today, Eurasia and Northern America are occupied by a handful of extant species and under this biogeographic scenario, the current occupation these Laurasian regions is the result of relatively recent events of colonization from the south. Likewise, several out-of-Australasia dispersal events are inferred into Southeastern Asia, India, and Eurasia, which reflect the presumed (relatively) recent colonization of these areas by species of *Plagiogyria*, *Culcita*, and *Gymnosphaera*.

Here I show that including fossil distribution information in biogeographical analyses of Cyatheales lead to substantially different results^36^. The inferred biogeographic history of a relatively recent occupation of Laurasian landmasses during the Paleocene is at odds with paleobotanical evidence. More specifically, the fossil records contrasts with the inferred Gondwanan origin of the Cyatheales and its constituent families (**figure 2a-b**). Non-overlapping fossil and extant geographic distributions are observed in several groups of ferns^16,20–26^, including several tree fern families. In particular, the fossil record places multiple Cyathealean lineages within Laurasia during the Jurassic and Cretaceous, and up to the Ecocene^18,27–30,42–44,83–86^, and some of the oldest known fossils (Late Jurassic) have been found in Eurasia and Northern America^29,42^. For instance, the Early Cretaceous *Kuylisporites mirabilis* and *Cyathea cranhamii* are the two oldest known representatives of the Cyatheaceae and were recovered from sites in Siberia^42^ and Canada^29^, respectively. Paradoxically, these fossils have been used to calibrate the age (stem or crown) of Cyatheaceae^38,56,77,87^ and produce the dated phylogenies underlying the biogeographical reconstructions supporting a Gondwanan ancestor^38^.

The presence of tree fern fossils at different points in time in regions that are currently uninhabited by extant members is evidence of a history (partly) hidden from biogeographic reconstructions; importantly, many Laurasian tree ferns fossils date back to at least the Late Cretaceous, at times when the crown diversification of most families was already underway. In this case, data and models of range evolution based on living tree ferns are limited and unable to account for the Laurasian legacy of the group. By bringing together paleontological and neontological information, I provide an alternative view on the biogeographic history of the Cyatheales where the most recent common ancestor of extant Cyatheales was probably distributed across Northern Pangea (later Laurasia) and then quickly spread into Southern Pangea (later Gondwana) during the Late Triassic–Early Jurassic; but with a disjunct distribution probably reflecting taphonomic biases (**figure 2c**). This is supported by the earliest tree fern fossils known from Gondwanan landmasses, which come from localities in Antarctica from the Middle to Lower Jurassic^88^. The early disjunct distribution of Cyatheales is consistent with paleobotanical evidence supporting a Gondwanan–Laurasian connection during the period^3,89,90^, in the form of a biotic corridor probably extending throughout ever wet climates around the Tethys sea^89,90^. Furthermore, recent evidence from the fossils *Paralophosoria*^91^ and *Eboracia*^27^, which have probable Cyathealean affinities, suggest the occupation of more meridional paleolatitudes (present-day Mexico) during this period.

It is likely that lineages of Cyatheales experienced major geographic range shifts and regional extinctions throughout their evolution associated with the paleogeography of continental drift^3,38,92^ and changes to global climate^3,93–96^. More specifically, I hypothesize that the disappearance of (most) tree ferns from Laurasian landmasses was likely triggered by changing climate during the Cenozoic, which began to shift into the drier and colder conditions that lead to the disappearance of tropical climates^19,94^. For instance, the Thyrsopteridaceae, now restricted to a single species in the Juan Fernández Island off the coast of South America^41,44^, is believed to have been more diverse and wide spread in the past, with fossils from the Cretaceous–Late Eocene uncovered in Japan^97^, Southeast Asia^44^, Australia^84,98^, and North America^99^.

The reconstruction using fossil data (*full*) supports probable secondary dispersals into Gondwana during the Late Jurassic–Cretaceous of the two most speciose families, the Cyatheaceae and the Dicksoniaceae. These results suggest an Eurasia–Southeastern Asia transition along the stem branch of the Cyatheaceae during the Late Jurassic–Early Cretaceous, temporarily coinciding with the inferred acceleration of diversification rates. Furthermore, this reconstruction supports an ancient vicariant event in *Cyathea* occurring in the Late to Early Cretaceous (ca. 100 Mya) associated with the final separation of Gondwana and Laurasia^3^. This event resulted in one lineage most probably distributed in Australasia and the other in [North America + Eurasia]. Subsequently, as continents drifted apart, *Cyathea* dispersed into South America via a northern route that differs from the more recent southern dispersal route (from Africa or Australasia) inferred for the other three genera in the Cyatheaceae. Coincidentally, the characteristically tropical palm family (Arecaceae) shows a similar biogeographic history as *Cyathea*^19,100^; the oldest know palm fossils have been discovered in Europe and North America^100^, which points to a Cretaceous origin of the group within the megathermal rain forests of Laurasia^19,101^ and a subsequent dispersal across Gondwanan landmasses during a period of profound climate and vegetation changes^94,102^.

### The fossil record of Cyatheales and taphonomic bias

Fossils provide direct evidence about past distributions and have the potential to shed new light on the biogeographic history of clades^11,13^. However, the inferences on the range evolution of Cyatheales are likely affected by unequal rates of preservation (and recovery) of fossils across different regions. Taphonomic bias is particularly relevant for the detailed assessment of regional occupancy and estimation of dispersal and extinction events. Thus, it is important to ask whether the absence of Cyatheales fossils in Gondwana during the Triassic–Lower Jurassic is a true reflection of their paleobiogeography. The present approach assumes that fossil sampling is representative of the paleodistribution of tree ferns and that the impacts of the regional heterogeneity in fossilization rates are negligible.

Even with an incomplete view of paleodistributions, the available evidence supports the undeniable past occupancy of Laurasia by members of the Cyatheales during the Late Triassic and Early Jurassic^27,103–109^, which extends into the Cenozoic. It is possible that the members of the Cyatheales initially inhabited Gondwana but failed to fossilize (or have not yet been recovered). However, recent fossil evidence from the Jurassic of present-day Mexico and China expand the know paleodistribution of Cyatheales across Laurasia during the Lower Jurassic^27,91^ and available evidence does not support the presence of Cyatheales in Gondwana prior to the Middle Jurassic, despite there being well studied localities in southern continents^89,90,110–112^. More specifically, Cyathealean tree ferns first appear in Gondwanan landmasses towards the Middle Jurassic^88^ and start to be more frequent into the Late Jurassic–Early Cretaceous^113–115^; palynological records provide evidence for the presence of tree ferns in South America until the Early Cretaceous^116^.

The taxonomy assignment of fossil species can mislead inferences on the paleodistribution of lineages; this is exemplified by the fossil genus *Coniopteris*, whose species appear towards the base of the tree fern phylogeny. Traditionally, members of *Coniopteris* have been placed within the tree ferns, but recently several species have been aligned with polypod ferns based on phylogenetic analyses^117^. Nevertheless, the *Coniopteris*-Polypodiales relationship shows minimal support as the key character supporting this relationship (the presence of a vertical and incomplete annulus) is missing for most species^117,118^. Furthermore, several *Coniopteris* species show the characteristic oblique (or nearly vertical) and complete annulus of the Cyatheales^117^ and, interestingly, these all come from Laurasian localities.

Phylogenetic uncertainty in the placement of fossils can also have a non-negligible impact on the inferred biogeography of Cyatheales. Although phylogenetic relationships can be inferred through the use of a total evidence approach, high-quality data are needed to make robust inferences^63^. This is a complex challenge because of the incomplete and fragmentary nature of fossil specimens, which is compounded in Cyatheales by the lack of comprehensive character matrices for extant members. In my analyses, the phylogenetic position of fossils is not informed by morphological data and is solely dependent on the FBD process and topological constraints. I attempted to minimize the effects of this lack of phylogenetic resolution by implementing a variant of the joint estimation of Landis et al.^73^; here the geographic distribution of extant and fossil species directly informs the model and favors tree topologies that are in harmony with biogeographic data.

## Conclusions

By combining paleontological and neontological distribution data for Cyatheales, I inferred a more complex biogeographical history than previously depicted based on the distribution of extant species alone. My analyses support the hypothesis that Cyatheales, particularly lineages outside Cyatheaceae, were more diverse and geographically more wide spread than at present times, spanning Gondwana and Laurasia during the Jurassic– Early Cretaceous. The present works shows than integrating fossils into event-based biogeographic analyses substantially alters inferences on the biogeographic history of Cyatheales; details on dispersal and extinction events might change when additional tree fern fossils are incorporated and their phylogenetic relationships are robustly inferred. The Cyatheales are witness to the impacts that geographically biased extinctions can have our capacity to infer ancestral geographic ranges and, very likely, other macroevolutionary trends. The fossil record is not without temporal, geographic, and phylogenetic bias, yet fossils alone hold information about past distributions and have the potential to inform models of range evolution and unveil hidden aspects of the biogeographic history of lineages.

## Acknowledgements

The Zenil Lab at the University of Kentucky, the Laboratorio de Evolución Molecular y Experimental at UNAM, and the Magallón Working Group at UNAM for lending of computational resources; Rosana Zenil-Ferguson and Michael Landis for sharing scripts to run the FBD and the DEC analyses in RevBayes; Carrie Tribble and Jaime Gasca Pineda for help setting up and troubleshooting RevBayes.

## Data and code availability

All data and code used to perform the biogeographical and macroevolutionary analyses are available at https://github.com/spiritu-santi/Biogeography.

