## supplementary for "Laurasian legacies in the Gondwanan tree fern order Cyatheales"

- This PDF file contains supplementary table S1 and figures S1–S3.
- Supplementary data can be found in the accompanying files.

Supplementary data 1. List of fossils specimens.

Supplementary data 2. DNA sequence alignment.

Supplementary data 3. Maximum likelihood phylogeny.

Supplementary data 4. Time-calibrated phylogeny including fossils.

Supplementary data 5. Time-calibrated phylogeny excluding fossils.

Supplementary data 6. BAMM control file.

Supplementary data 7–8. BAMM sampling fraction files.

**Supplementary table S1. Divergence time estimates for families and genera of tree ferns (Cyatheales).** Median age estimates, 95% high posterior density age estimates (95%HPD), and posterior probability (PP) derived from the time calibrated phylogeny under the Fossilized Birth-Death process. Ages are given for all families and genera.

|  | Crown ages | | | |  | Stem ages | | | |
| --- | --- | --- | --- | --- | --- | --- | --- | --- | --- |
|  | median | 95%HPD | | PP |  | median | 95%HPD | | PP |
| Cyatheaceae | 121.84 | 104.77 | 139.42 | 1.00 |  | 178.89 | 159.64 | 197.76 | 0.72 |
| *Cyathea* | 70.65 | 53.39 | 91.62 | 1.00 |  | 111.37 | 100.00 | 123.80 | 1.00 |
| *Alsophila* | 56.60 | 38.71 | 76.48 | 1.00 |  | 77.38 | 50.61 | 110.17 | 0.74 |
| *Gymnosphaera* | 51.45 | 28.66 | 79.87 | 1.00 |  | 77.38 | 50.61 | 110.17 | 0.74 |
| *Sphaeropteris* | 81.68 | 52.85 | 115.51 | 1.00 |  | 121.84 | 104.77 | 139.42 | 1.00 |
| Dicksoniaceae | 138.05 | 124.26 | 155.50 | 1.00 |  | 166.67 | 144.72 | 189.46 | 0.80 |
| *Dicksonia* | 31.15 | 16.87 | 52.27 | 1.00 |  | 122.67 | 100.84 | 145.57 | 1.00 |
| *Calochlaena* | 24.20 | 8.36 | 42.35 | 1.00 |  | 122.67 | 100.84 | 145.57 | 1.00 |
| *Lophosoria* | 2.98 | 0.06 | 9.56 | 1.00 |  | 138.05 | 124.26 | 155.50 | 1.00 |
| Cibotiaceae* | 23.44 | 8.86 | 44.98 | 1.00 |  | 166.67 | 144.72 | 189.46 | 0.80 |
| Metaxyaceae* | 15.94 | 6.63 | 27.17 | 1.00 |  | 188.08 | 169.58 | 207.86 | 1.00 |
| Plagiogyriaceae* | 39.87 | 20.28 | 61.90 | 1.00 |  | 74.85 | 48.32 | 107.65 | 1.00 |
| Culcitaceae* | 19.22 | 3.15 | 42.98 | 1.00 |  | 74.85 | 48.32 | 107.65 | 1.00 |
| Loxomataceae** | 32.86 | 9.42 | 62.50 | 1.00 |  | 140.09 | 118.36 | 167.98 | 1.00 |
| Thyrsopteridaceae | - | - | - | - |  | 179.04 | 143.80 | 214.16 | 0.85 |

*Stem and crown ages for Cibotium, Metaxya, Plagiogyria, and Culcitaceae are the same as those of the corresponding families.

** Stem ages for Loxsoma and Loxsomopsis are the same as the crown age of the family.

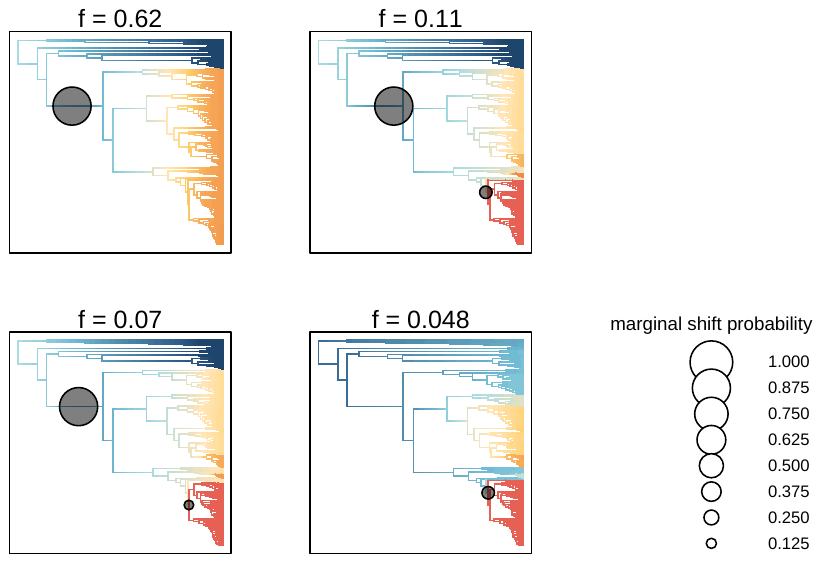

**Supplementary figure S1. Four best shift configurations obtained in the BAMM using a global sampling fraction.**

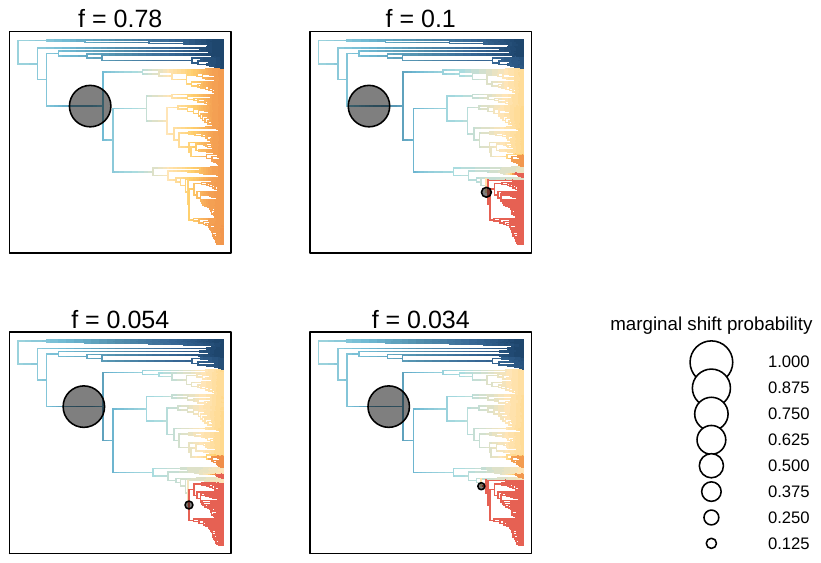

**Supplementary figure S2. Four best shift configurations obtained in the BAMM using a per genus sampling fraction.**

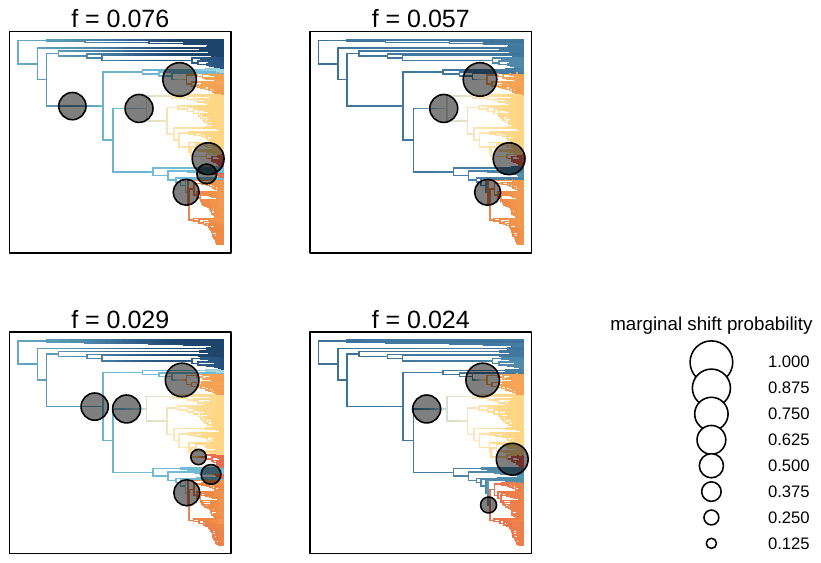

**Supplementary figure S3. Four best shift configurations obtained in the BAMM using a per lineage sampling fraction.**

1. correspondance to
